# Neuronal resonance in the theta (4-10 Hz) frequency range is modulated by dynamic changes in the input resistance

**DOI:** 10.1101/224253

**Authors:** Jorge Vera, Ulises Pereira, Bryan Reynaert, Juan Bacigalupo, Magdalena Sanhueza

## Abstract

Most neurons of the mammalian brain display intrinsic resonance with frequency selectivity (*f_R_*) for inputs within the theta-range (4-10 Hz). Variations in network oscillation along this range depend on the animal behavior; however, whether neurons can dynamically tune their *f_R_* has not been addressed. Using slice electrophysiology, dynamic clamping and computer modeling we studied three types of cortical neurons, finding that the input resistance (*R_in_*) inversely sets *f_R_* into the theta range, following a power law. We demonstrate that physiological changes in *R_in_* modulate *f_R_* and response phase, serving as a mechanism that instantaneously tunes oscillatory responses. Moreover, these modulations are translated into spiking regimes, modifying spike frequency and timing. Since synaptic inputs reduce *R_in_*, this modulation provides a mean for adjusting the frequency and timing of firing of individual neurons in interplay with the network fluctuations. This might be a widespread property among resonant neurons.

## Introduction

Theta frequency oscillations (4-10 Hz) are observed in different brain regions associated to active sensory sampling, navigation and learning processes (Kay, 2005; Lisman & Buzsáki, 2008; Mizuseki *et al*., 2009; Ranade *et al*., 2013). These oscillations reflect synchronized neuronal activity spreading across different networks and support a neural code based on the frequency and timing of neural firing (Lisman and Jensen, 2013; Wilson et al., 2015).

Theta activity is displayed by several brain regions and varies with time according to the behavioral state of the animal, increasing or decreasing its frequency, depending on the animal behavior (Kay, 2005; Geisler *et al*., 2007; Ranade *et al*., 2013; Bender *et al*., 2015). This observation is in accordance to the emergent idea that neurons can modify their responsiveness relative to the network state (Destexhe *et al*., 2003; Ratté *et al*., 2013). For instance, neurons can switch their behavior to favor or hamper neural synchrony (Prescott *et al*., 2008; Kirst *et al*., 2017; Vera *et al*., 2017). Thus, it is expected that variations in the frequency of the network oscillations would demand flexibility of the intrinsic activity of the neurons, in order to follow such ongoing dynamic changes.

Many cortical neurons resonate, displaying an enhanced voltage response in the theta range, strengthening input signal processing and transmission at those frequencies (Hutcheon *et al*., 1996a; Ulrich, 2002; Hu *et al*., 2002; Erchova *et al*., 2004; Vera *et al*., 2014). This intrinsic property relies on the presence of the hyperpolarization-activated cationic current (*I*_h_) (Biel *et al*., 2009) and/or the muscarine-sensitive current (*I_M_*) (Shah *et al*., 2002), that selectively reduce the low frequency (<10 Hz) voltage responses due to their slow activation/deactivation kinetics (Hutcheon and Yarom, 2000; Izhikevich, 2002).

This property can be phenomenologically described by an inductor (Mauro, 1961) that reduces not only the impedance, but also the phase-lag, which is the frequency-dependent delay of the voltage response with respect to an oscillatory stimulus. The phase-lag approaches zero at particular frequencies in the theta range (Narayanan & Johnston, 2008; Rotstein, 2014), thus functioning as a mechanism to synchronize synaptic inputs (Vaidya & Johnston, 2013).

The frequency at which the maximal impedance is observed is called resonance frequency (*f_R_*), and is set by a combination of passive and active intrinsic membrane properties (Hutcheon & Yarom, 2000). *In vitro* experiments have shown that most resonant neurons in the mammalian brain display *f_R_*s between 2 and 10 Hz: isocortex (Gutfreund *et al*., 1995; Hutcheon *et al*., 1996a), thalamus (Puil *et al*., 1994), hippocampus (Hu *et al*., 2002), entorhinal cortex (Erchova *et al*., 2004; Giocomo *et al*., 2007), amygdala (Pape & Driesang, 1998; Vera *et al*., 2014) and olfactory bulb (Hu *et al*., 2016). However, it is not clear whether the differences in *f_R_* represent variations in the resonant mechanism or are due to different experimental conditions. Previous investigations have shown that *f_R_* can increase due to a rise in the amplitude of the *I_h_* current, caused by the induction of long term synaptic potentiation (Narayanan & Johnston, 2007). A dynamic change in *f_R_* is also observed during membrane depolarization (Gutfreund et al., 1995; Hutcheon et al., 1996a and b), however, this process has not been observed in all resonant cells (Erchova *et al*., 2004).

One relevant parameter involved in setting *f_R_* is the passive membrane resistance (*R_m_*, the reciprocal of the leak conductance), that together with the membrane capacitance (*C*_m_) sets the cut-off frequency of the low-pass filter that determines the frequency response (*fc_m_*=1/2πτ_m_, where τ*_m_=C_m_*R_m_*)(Hutcheon & Yarom, 2000). The presence of *I_h_* or *I_M_* endows resonant neurons with a high-pass filter whose cut-off frequency depends on the kinetic properties of the conductances (Hutcheon & Yarom, 2000). The combination of these two filters produces a bell-shape impedance profile that defines the frequency selectivity into the theta range, with *f_R_* corresponding to the peak of the resulting band-pass filter (Hutcheon & Yarom, 2000). Thus, it can be expected that changes in *R_m_* will modify the steepness of the low-pass filter, therefore shifting *f_R_*. Moreover, changes in *R_m_* also modify the membrane time constant, drastically altering the timing of neuronal responses (Destexhe *et al*., 2003; Broicher *et al*., 2012) and probably modifying the phase-lag at theta frequencies.

Measurement of *R_m_* is hampered by the presence of *I_h_* or *I_M_,* whose voltage-and time-dependent activation deviate the voltage behavior from the purely passive response. Instead, we used a current pulse to measure what we refer to as input resistance (*R_in_*) of a neuron, which at frequencies near zero is directly related to the total membrane conductance at steady state.

We predict that neurons with different *R_in_*s would display different *f_R_*s. Moreover, sources of *R_in_* variations such as voltage-dependent and independent, as well as synaptic conductances could modify it, generating variations in the *f_R_* and the kinetics of the responses. Whether this *R_in_* dependence is relevant in setting *f_R_* and the phase-lag of the responses in theta range has not been evaluated.

We characterized how *f_R_* and the phase-lag depend on *R_in_* in three different cortical neurons types that share the same resonant mechanism, but differ in their *R_in_* range: stellate neurons from layer II of the entorhinal cortex (SL)(Erchova *et al*., 2004), CA1 pyramidal neurons from the hippocampus (HP)(Hu *et al*., 2002) and layer II resonant neurons of the cortical amygdala (AM)(Vera *et al*., 2014). To evaluate frequency selectivity upon changes in *R_in_,* we considered, i) the natural differences in *R_in_* among cell types, ii) changes in *R_in_* due to activation/deactivation of voltage-sensitive conductances upon membrane depolarization, and iii) changes in *R_in_* caused by introducing virtual passive conductances by means of dynamic clamp (Dorval *et al*., 2001).

We found that neurons exhibit a frequency selectivity that is determined by *R_in_* according to a power law distribution, setting *f_R_* values within the theta range. Moreover, variation in *R_in_* alters the phase-lag curves at frequencies below 15 Hz. The relationship between *f_R_* and *R_in_* is robust for different cortical neurons, as well as for *R_in_* variations within individual cells. This modulation is observed in both subthreshold and spiking regimes, modifying spike frequency and timing.

## Methods

### Ethical approval

Animal care and experimental procedures were approved by the Bio-Ethical Committee of the Faculty of Sciences, University of Chile, according to the ethical rules of the Biosafety Policy Manual of the National Fund for Development of Science and Technology (FONDECYT).

### Slice preparation

Male Sprague Dawley rats (18 to 30 days-old) were deeply anesthetized with isoflurane, decapitated and their brain was rapidly removed and transferred to an ice-cold dissection solution containing (in mM): 206 sucrose, 2.8 KCl, 1 MgCl_2_ MgSO_4_, 1 CaCl_2_, 26 NaHCO_3_, 1.12 NaH_2_PO_4_, 10 glucose and 0.4 ascorbic acid (equilibrated with 95% O_2_ and 5% CO_2_), pH 7.3. Slices (400 μm) containing the region of interest were obtained with a vibratome (Vibratome Sectioning System 102, Pelco) using standard disposable stainless steel razor blades (http://www.personna.com.) For the anterior nucleus of the cortical amygdala we used coronal slices targeting the region between bregma -2.2 and -3.3 (Sanhueza & Bacigalupo, 2005; Vera *et al*., 2014). For hippocampal neurons we used septotemporal slices targeting the dorsal hippocampus (Vera *et al*., 2017), and for stellate cell we used horizontal slices to target the dorsomedial entorhinal cortex(Giocomo *et al*., 2007). Slices were placed in a holding chamber with standard artificial cerebro-spinal fluid (ACSF) and were left to recover at least 1 h at 30°C before using them for recording.

### Electrophysiological recordings

Whole cell patch-clamp recordings were conducted under visual guidance by an upright microscope (Nikon Eclipse E600FN) equipped with DIC optics. Patch pipettes (3.5–4.5 MΩmΩ) were fabricated of borosilicate glass (0.8 - 1.10 x 100 mm; Kimble Glass Inc) using a horizontal puller (Flaming/Brown P-97, Sutter Instrument Co). Current-clamp recordings were made with an EPC-10 patch-clamp amplifier (Heka, Heidelberg, Germany), data were filtered at 5 kHz and acquired at 40 kHz using the Patch Master software from Heka. All experiments were performed in the presence of 10 μM CNQX and 100 μM PTX, to block AMPA-R and GABAa-R mediated currents.

### Recording solutions (in mM)

Artificial cerebro-spinal solution (ACSF) contained (in mM): 124 NaCl, 2.8 KCl, 1.25 NaH_2_PO_4_, 26 NaHCO_3_, 10 Glucose, 2 MgCl_2_, 2 CaCl_2_ and 0.4 ascorbic acid (equilibrated with 95% O_2_ and 5% CO_2_), pH 7.3 and 290 mOsm.

Internal pipette solution contained (in mM): 123 K-Gluconate, 10 KCl, 4 Glucose, 1 EGTA, 10 HEPES, 2 Na_2_ATP, 0.2 Na_3_GTP, 10 phosphocreatine-Na, 1 MgCl_2_, 0.1 CaCl_2_, pH 7.35 and 290 mOsm. This composition was based on previous works reporting stability of intrinsic excitability parameters (Xu *et al*., 2005; Kaczorowski *et al*., 2011), and was corroborated in investigations related to intrinsic frequency preference (Vera *et al*., 2014, 2017). We measured the liquid junction potential (LJP) between pipette solution and ACSF (~13 mV) according to the procedure described by Neher (Neher, 1992) and used this to correct the data in offline analyses.

### ZAP stimulation and data analysis

The ZAP (impedance amplitude profile) stimulus consisted in a pseudo-sinusoidal current of constant amplitude and linearly increasing frequency from 0 to 20 Hz in 10 s (ZAP stimuli). The protocol was repeated 8 to 10 times in every neuron and the membrane voltage waves were averaged for the impedance analysis. The impedance profile (*Z(f)*) and phase-lag (Φ*(f)*) curves were obtained from the ratio of the Fast Fourier Transforms (FFT) of the output (voltage) and input (current) waves (*Z(f)=FFT[V(t)]/FFT[I(t)*]). The impedance is a complex quantity (*Z(f)* = *Z_Real_* + *iZ_Imaginary_*), where the real part (*Z_Real_*) is the resistance and the imaginary part (*Z_Imaginary_*), the reactance. For each given frequency, the complex impedance can be plotted as a vector whose magnitude and phase (Φ*(f)*; angle with the real axis) are respectively given by the following expressions:

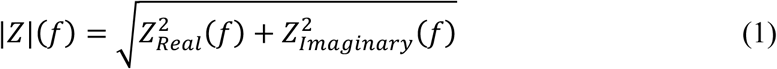

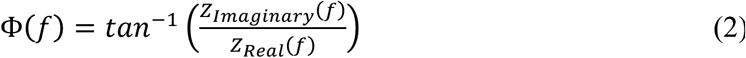

Throughout the text the term *impedance* is used to refer to the magnitude of the impedance vector, unless otherwise stated. The impedance phase corresponds to the phase shift of the voltage wave relative to the current wave. Frequencies below 0.2 Hz were not plotted in the impedance and phase profiles graphs, to avoid low frequency distortions.

### Quantification of the resonance

Resonance is defined as the band-pass filter property of the impedance profile (Hutcheon & Yarom, 2000). The resonance strength is usually quantified as the ratio between the maximal impedance (i.e. the impedance at the resonance frequency, (|*Z(f_R_)*|) and the impedance at 0.5 Hz (|*Z(0.5)*|). This ratio is termed *Q*, and indicates the sharpness of the impedance curve around the resonance frequency. For a more precise determination of *Q*, the experimental data were fitted with a polynomial curve between 0.2 and 15 Hz and the peak value was calculated.

### Dynamic clamp experiments

For the dynamic-clamp experiments, the current-clamp amplifier was driven by an analog signal from a dual core desktop computer running the Real-Time Experimental Interface, RTXI (Dorval et al., 2001), using an update frequency of 25 KHz.

The somatic conductance was altered by simulating a constant conductance via the dynamic clamp. The leak current was computed according to the following equation:

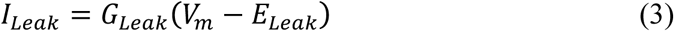

where *G_Leak_* is the evaluated leak conductance, *V_m_* is the online recorded membrane potential and *E_Leak_* is the reversal potential (set to -70 mV). To evaluate a decrease in *R_in_, G_Leak_* > 0; for an increase in Rin, *G_Leak_* < 0.

We show dynamic currents (*I_DynC_*) as the external current that was injected to neurons, following the standard convention that positive currents are depolarizing and negative currents are hyperpolarizing. Therefore,

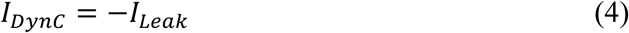

### Computer simulations

We developed a minimal point process and conductance-based model following the Hodgkin-Huxley formalism (Hodgkin & Huxley, 1952). The model included a membrane capacitance (*C_m_*), a passive leak current (*I_Leak_*), the hyperpolarization-activated *h* current, *I_h_* (Richardson et al., 2003) and an external stimulating current, *I_ZAP_* The model did not include a firing mechanism (fast sodium and potassium conductances).

The equation describing the evolution of the membrane potential (*V*) in time is

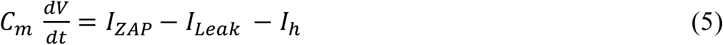

where *C_m_* is the membrane capacitance and *I_ZAP_* is the externally applied current. The intrinsic ionic currents in Eq. 1are described by the following equations:

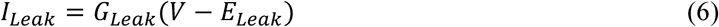

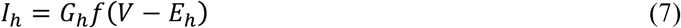

where *G_Leak_* and *G_h_* are the maximal conductances of the corresponding currents and *E_Leak_* (-65 mV) and *E_h_* (-40 mV) are the reversal potentials of *I_Leak_* and *I_h_*, respectively. *f* is the state variable of *I_h_* (Spain *et al*., 1987). For simplicity, in our minimal model we only used the fast component of *I_h_*, which is related to theta resonance and accounts for ~80% of the total current (Spain *et al*., 1987).

The dynamics of the state variable *f* is ruled by the following equation:

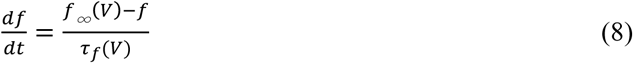

where *τ*_*f*_ is the time constant and *f*_∞_ is the steady-state value of *f*, calculated according to:

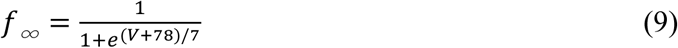

*τ_f_* was set to 0.05 s, similar to the fast component described by Spain et al. (Spain *et al*., 1987), to match the exact *f_R_* range observed in our experiments.

We reproduced each cell type using the membrane capacitance estimated from voltage-clamp recordings (50 ms, 5 mV pulse) and the maximal conductances were tuned to obtain similar voltage responses at -80 mV (Table 1, Fig. 5). When exploring the effect of *G_Leak_,* we varied its value from 1 to 80 nS.

Simulations were performed using Igor Pro 6.37 software with an integration time step of 10 μs. The code for reproducing the computer simulations described in this paper is available upon request to authors.

### Statistical analysis

The statistical analysis was performed in GraphPad Prism 6.07 (GraphPad Software, Inc, USA). Group data is presented as the mean ± standard error. We used different parametric tests for data with normal distribution (as *Z_Max_,* resting potential or *R_in_*). When the data structure was a single variable (*Z_Max_*, *R_in_*, *f_R_*) measured at the same membrane potential and compared between different cell types (Fig. 1) or was coming from the same cell type but comparing different membrane potentials (Fig. 2), we used one-way ANOVA with matched samples and Tukey’s multiple comparison tests. When the comparison of variables was between two conditions in the same cell (Fig. 3), we used paired Student’s t-test. To evaluate differences in firing probability as a function of stimulation frequency after reducing *R_in_* (Fig. 4), we used two-way repeated measures ANOVA with Holm-Šídák’s multiple comparison tests. The statistical significance of the linear relation between *f_R_* and *R_in_* (Figs. 1 and 2) was evaluated with a Student test. Statistical tests were two-tailed and we used α=0.05 as critic value.

**Table 1.**
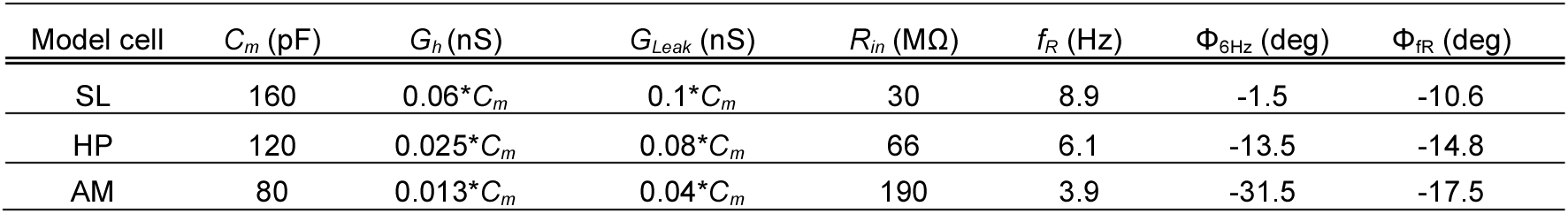
Parameters used in (*C*_m_, *G*_h_ and *G*_Leak_) and obtained from (*R*_in_, *f_R_*, Φ_6Hz_ and Φ_fR_) simulated neurons.

## Results

### The preferred frequency and phase-lag of cortical neurons are correlated with the input resistance

To investigate the connection of *R_in_* with *f_R_* and the phase-lag, we conducted whole cell recordings in three well described types of cortical neurons that share the same subthreshold resonant mechanism: entorhinal cortex layer II stellate (SL) neurons (Erchova *et al*., 2004), hippocampal CA1 pyramidal neurons (HP) (Hu *et al*., 2002) and cortical amygdala layer II resonant neurons (Vera *et al*., 2014). Figure 1a shows the voltage responses of representative neurons of each type to hyperpolarizing and depolarizing square current pulses. The responses to both stimuli polarities start with a sag at the onset of the pulse, produced by the slow (tens of milliseconds) activation or deactivation of the inward *I_h_,* respectively (Biel *et al*., 2009). At the end of the pulse, there is a transient rebound with similar kinetics (Biel *et al*., 2009). Further depolarization induced trains of action potentials with some frequency accommodation, presumably due to *I_h_* deactivation and the activation of the muscarine sensitive potassium current (*I*_M_) (Lawrence *et al*., 2006). These similarities in voltage responses and firing pattern corroborate that these neurons possess similar electrophysiological behaviors (Hu *et al*., 2002; Erchova *et al*., 2004; Vera *et al*., 2014).

*R_in_* was measured with 250 ms hyperpolarizing pulses adjusted to produce a ~5 mV maximal deflection from -80 mV. *R_in_* values of the three types of neurons ranged from 10 to 400 MΩ, where SL neurons are more abundant in the lower (mean = ~26 MΩ), HPs in the intermediate (mean = ~65 MΩ) and AMs in the highest *R_in_* sub-ranges (mean = ~200 MΩ; Fig. 1b, Table 2). Our observations corroborate that the three cell types are suited for investigating how *R_in_* influences their *f_R_* and phase-lag.

To quantify and compare the resonant properties of these neurons we recorded their voltage responses to oscillatory stimulation. We applied ZAP (impedance amplitude profile) stimuli (Puil *et al*., 1986), consisting of a 10 s oscillatory current of constant amplitude and linearly incremental frequencies, from 0 to 20 Hz. The membrane potential was manually adjusted to ~-80 mV in order to set a similar level of activation of *I_h_* and the stimulus amplitude was adjusted to produce a comparable voltage deflection of ~5 mV peak to peak at its onset (ranging from 10 to 50 pA). As expected (Hutcheon & Yarom, 2000), the voltage response in all tested cells displayed a maximal amplitude in the theta range (arrows) and the low and high frequencies of their responses were attenuated (Fig. 1c).

**Figure 1.**
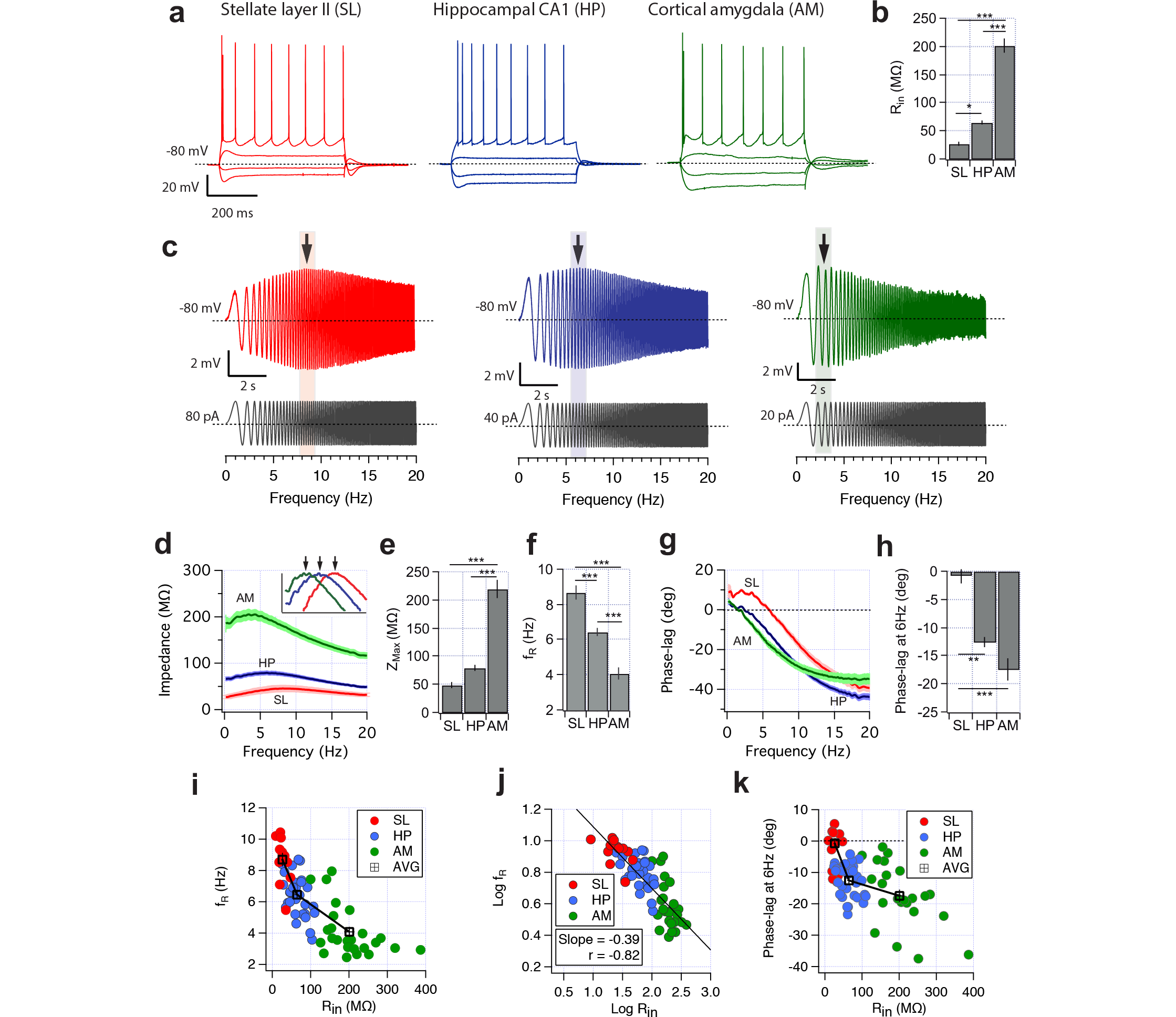
Frequency preference of resonant cell types (peak impedance and --lag) correlates with their Rin. Impedance analysis performed in stellate neurons from layer II entorhinal cortex (SL, n=12), pyramidal cells from CA1 hippocampus (HP, n=31) and resonant neurons from the anterior cortical amygdala (AM, n=25). **a** Representative voltage response of SL (red), HP (blue) and AM (red) neurons. Current pulses were -150,-50,+100,+150 pA for SL, -200,-100,+50,+100 pA for HP and -100, -50, +50, +100 pA for AM. **b** Input resistance of SL, HP and AM neurons measured at -80 mV with a hyperpolarizing squared pulse. **c** Representative voltage responses of each cell type stimulated with the ZAP protocol (grey traces, see Methods). Peak voltage responses are shown with arrows. **d** Impedance profile curves from each cell types (dark and light colors depict mean and SEM, respectively); the inset shows normalized curves from 0.8 to 1 (y axes) and from 0 to 14 Hz (x axes). Arrows shows the peak of each curve. **e** Population peak impedance measured from impedance curves. **f** Population *f_R_* measured from impedance profile curves. **g** Phase-lag curves, dark and light colors depict mean and SEM, respectively. **h** Quantification of phase-lag at 6 Hz (Φ_6Hz_). **i** Scatter plots of *f_R_* vs. *R_in_* with data from all recorded neurons (n=68); **j** Scatter plot of *f_R_* vs. *R_in_* transformed to log(*f_R_*) vs. log(*R*_in_). **k** Scatter plot of Φ_6Hz_ vs. *R_in_* One way ANOVA, * = P < 0.05,** = P < 0.01 and *** = P < 0.001.

**Table 2.** Electrophysiological parameters for stellate (SL), hippocampal (HP) and cortical amygdala (AM) neurons characterized at hyperpolarized potential. Mean ± SEM. Pairs for statistical comparison: 1 SL-HP, 2 SL-AM and 3 HP-AM. * P<0.05, **P<0.01, ***P<0.001, ****P< 0.0001.

| | $V_m$ (mV) | $Z_{Max}$ (MΩ) | $f_R$ (Hz) | $Q$ | $\Phi_{6Hz}$ (deg) | $\Phi_{fR}$ (deg) | $R_{in}$ (MΩ) | $n$ |
| --- | --- | --- | --- | --- | --- | --- | --- | --- |
| SL | $-79.6 \pm 0.4$ | $48 \pm 6.1$ | $8.7 \pm 0.4$ | $1.62 \pm 0.05$ | $-0.7 \pm 1.5$ | $-12.4 \pm 0.6$ | $26.5 \pm 3.2$ | 12 |
| HP | $-80.0 \pm 0.6$ | $78 \pm 5.0$ | $6.5 \pm 0.3$ | $1.23 \pm 0.02$ | $-12.6 \pm 0.1$ | $-13.9 \pm 0.6$ | $64.8 \pm 4.4$ | 31 |
| AM | $-80.6 \pm 0.3$ | $220 \pm 16.2$ | $4.1 \pm 0.3$ | $1.15 \pm 0.02$ | $-17.5 \pm 0.8$ | $-8.4 \pm 0.7$ | $200.9 \pm 13.4$ | 25 |
| Statistics | ns | 1(ns), 2****, 3**** | 1****, 2****, 3**** | 1****, 2****, 3(ns) | 1****, 2****, 3* | 1(ns), 2**, 3**** | 1*, 2****, 3**** |  |

The average impedance profiles of the three groups of neurons were clearly different. Their characteristic shapes resemble band-pass filters with peaks in the theta range (Fig. 1d), but their average maximal impedances (*Z_Max_*) and *f_R_*s were obviously dissimilar: the average *Z_Max_* was ~50 MΩ for SL, ~80 MΩ for HP and ~220 MΩ for AM neurons (Table 2, Fig. 1e) and their respective impedance curves are displaced in the frequency range, as can be better appreciated in the normalized impedance profiles (Fig. 1d, inset). The *f_R_*s derived from the impedance curves was higher in SL (8.7 ± 0.4 Hz, n = 12) compared to HP (6.5 ± 0.3 Hz; n = 31) and AM neurons (4.1 ± 0.3 Hz, n = 25; P < 0.001 for all paired comparisons; Table 2, Fig. 1f;). The resonance strength among the different cell populations, evaluated using the *Q* value (see Methods), was higher in the SL neurons, than the HP and the AM neurons (Table 2).

The time dependence of the voltage response was analyzed with phase-lag curves that quantify the delay of the output voltage wave relative to the current stimulus as a function of frequency. The negative phase-lag values reflect a delay of the voltage response and the mean of the positive values an advance, which may provide important insights of the temporal processing of oscillatory stimuli (Rotstein, 2014). The analysis of phase-lag curves showed that the response is faster (i.e., shorter lag) at frequencies within and below the theta range (0-8 Hz) for SL in comparison to HP and AM neurons (Fig. 1g). We quantified the phase-lag of each cell at two relevant frequencies: at the middle of theta range (6 Hz, Φ_6Hz_) and at the *f_R_* of each particular neuron (Φ_fR_). The Φ_6Hz_ values were markedly different between the three groups: -0.7 ± 1.5 deg, -12.5 ± 1.0 deg and -17.5 ± 2.0 deg for SL, HP and AM neurons, respectively (Fig. 1h, Table 2, P<0.001 in all paired comparisons). By transforming these lag values from degrees to time (where 360 deg are 166.7 ms in a single oscillation at 6 Hz) we obtain a phase-lag of -0.3 ms, -5.8 ms and -8.1 ms, for SL, HP and AM neurons, respectively (P<0.05 for each pair, see values in Table 2), presenting a difference in the temporal response that might be relevant for spike timing and coding in these cell types. When phase-lag is compared at the *f_R_* of each neuron the three cell types display less variability than at 6 Hz, with values grouped in the narrow range of 8-14 deg. The quantification shows similar phase-lags in SL and HP cells, with a reduced lag in AM neurons (-12.4 ± 0.6 deg, -13.9 ± 0.6 deg and -8.4 ± 0.7 deg for SL, HP and AM neurons, respectively. P < 0.001, 1 way ANOVA; see paired statistics in Table 2).

For a more detailed examination of the *f_R_* and *R_in_* relationship, we built a scatter plot with the values from all recorded cells. We found that *f_R_* increased as *R_in_* decreased, exhibiting a negative correlation. Low-*R*_in_ cells show the highest *f_R_s* and the lowest *f_R_*s were observed in the high-*R*_in_ cells (Fig. 1i). The *R_in_* range covered by the three cell types is associated with a continuum variation of *f_R_s* from about 2 to 10 Hz, matching the theta range. Since the distribution of the data in the scatter plot seems to represent a multiplicative relationship between *f_R_* and *R_in_,* we transformed the data to logarithmic values (Kass *et al*., 2014). The scatter plot of log(*f_R_*)-log(*R*_*in*_) pairs yields a linear relation characterized by a Pearson’s correlation coefficient of -0.82 and a slope of -0.39 (n = 68, P < 0.001), suggesting a power law between *f_R_* and *R_in_* (Fig. 1j).

We also explored the correlation between Φ_6Hz_ and *R_in_* pooling the data from all cells in a scatter plot. Φ_6Hz_ is also correlated with *R_in_,* dominating the values around zero lag for lower *R_in_* cells, while for higher *R_in_* the phase values can reach ~-40 deg (Fig. 1k).

This comparative analysis revealed a co-variation between the resonant parameters (*f_R_* and Φ_6Hz_) and the magnitude of *R_in_* at -80 mV. It is noteworthy that, despite all the factors that could influence *R_in_* in neurons from distinct brain regions (like the magnitude and distribution of membrane conductances, dendritic and somatic morphologies, etc.), it is still possible to find such a high degree of correlation with the resonance. This result shows that the *R_in_* of the neurons is associated with a particular setting of resonance, producing cell-type specific *f_R_*s along the theta range and different phase-lag, even though all cells tested possess the same active resonant mechanism.

### Neurons modify the frequency preference and phase-lag during depolarization-driven increase in *R_in_*

To investigate to what extent the variation in *R_in_* was associated with modifications in *f_R_* and phase-lag, we changed *R_in_* by manipulating the activation levels of the voltage-sensitive conductances with current injection, an operation that does not modify the cell passive properties. Steady-state depolarization of the cortical neurons from -80 mV to near threshold potentials produced a reduction of *I_h_* due to deactivation (Hutcheon *et al*., 1996*a*; Hu *et al*., 2002; Biel *et al*., 2009) and the activation of the persistent Na^+^ current, *I_NaP_,* above -70 mV (Crill, 1996; Vera *et al*., 2014, 2017; Ceballos *et al*., 2017). Both modifications produce a voltage-dependent rise in *R_in_* that has been observed previously in resonant neurons (Gutfreund *et al*., 1995; Hutcheon *et al*., 1996a), and characterized in detail more recently (Surges *et al*., 2004; Economo *et al*., 2014; Yamada-hanff & Bean, 2015; Ceballos *et al*., 2017); we will refer to it as “depolarization-driven increase in R_in_”. We manually adjusted the membrane potential to -80, -70 and -60 mV by current injection and applied the ZAP stimuli at each potential, exploring *R_in_* and resonance in SL and AM neurons. We did not include HP neurons in this experiment because they present a U-shaped voltage dependence of the resonance, lacking frequency preference between -70 and -65 mV (Hu *et al*., 2002; Vera *et al*., 2017).

The shape of the voltage responses to ZAP stimuli changed with depolarization, revealing an increased filtering at high frequencies and an early peak of the voltage response, manifesting a reduction in the *f_R_* (Fig. 2a). *R_in_* increased with depolarization from 26 to 65 MΩ in SL neurons and from 167 to 290 MΩ in AM neurons, with an equivalent increase in peak impedance (traces in Fig. 2b,f, and values in Table 3). This wide variation in *R_in_* allowed a good comparison of this variable with the *f_R_* and phase-lag. We confirmed that the increase in *R_in_* was accompanied by a reduction in *f_R_* from 8.7 to 6 Hz in SL neurons (Fig. 2c), and from 3.6 to 2.7 Hz in AM neurons (Fig. 2g, Table 3). The depolarization was also accompanied by a modification in the phase-lag profiles of both cell types, shifting the curves downward and leftward, with the consequent increment in lag as a function of frequency (Fig. 2d, h). The overall change in the phase-lag curves produced a considerable increase of Φ_6Hz_ in SL and AM neurons (-0.75 to -25.8 deg and -17.0 to -34.8 deg, respectively; Fig. 2e, i, Table 3). However, when quantified at *f_R_*, the phase-lag showed a smaller range of variations under depolarization, increasing Φ_fR_ from -12 to -20 deg inSL neurons (P<0.001, 1 way ANOVA), while in AM neurons it stayed near -8 deg without significant changes (P=0.16, 1 way ANOVA; see details in Table 3).

**figure 2.**
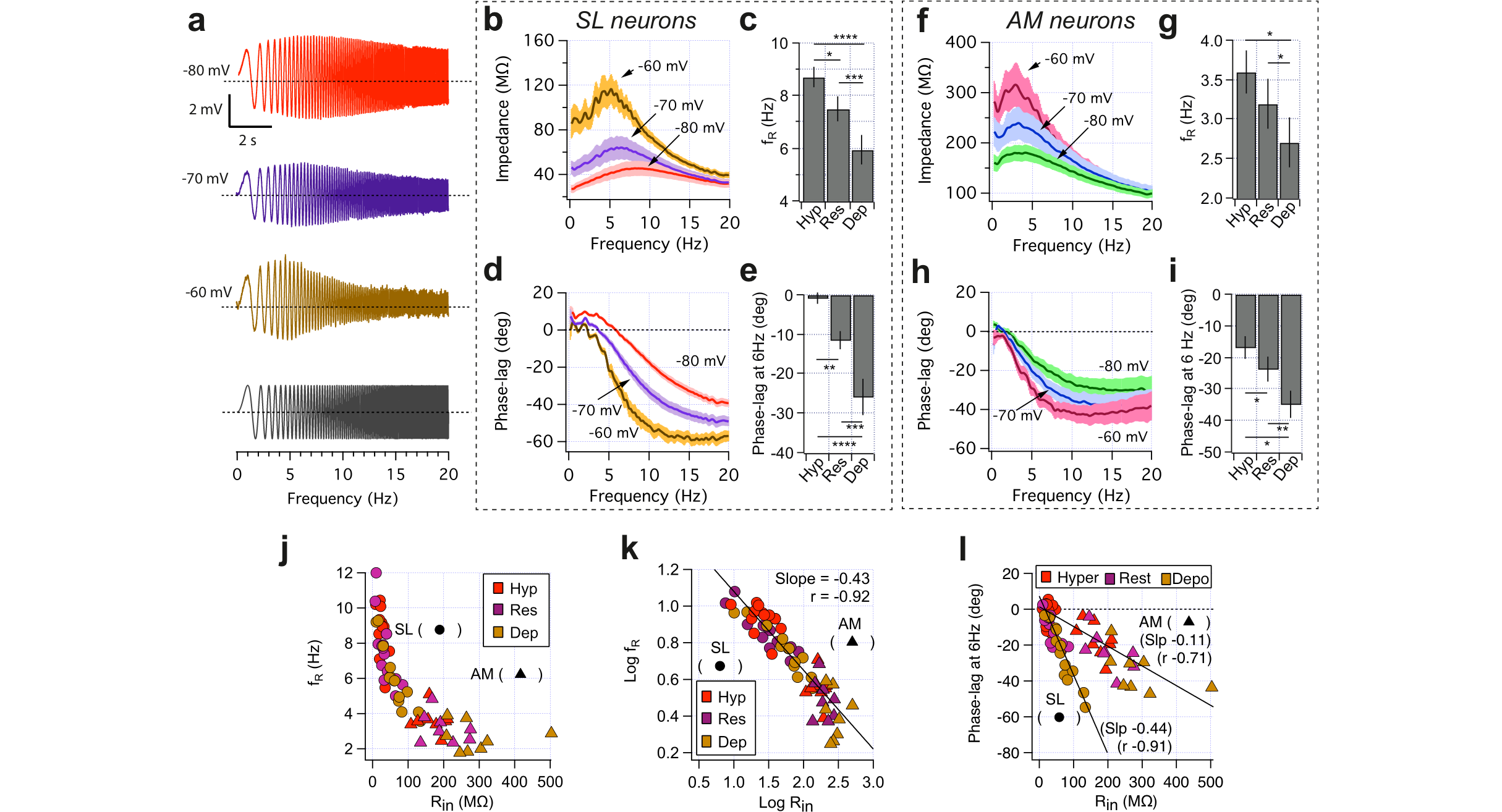
Depolarization-driven increase in *R_in_* correlates with reductions in *f_R_* and increments in phase-lag. Paired measurement of resonant properties in SL (n=11) and AM (n=8) neurons recorded at hyperpolarized (Hyp), resting (Res) and depolarized (Dep) membrane potential (see Table 3), to evaluate the effect of depolarization. **a** Representative voltage response of a SL neuron to a ZAP stimulus. The amplitude of the sinusoidal current (gray trace) was adjusted in order to maintain voltage oscillations near 5 mV (in this example, 40, 20 and 5 pA for -80, -70 and -60 mV, respectively). **b** Impedance profile curves of SL neurons (mean ± SEM). **c** Quantification of *f_R_* as a function of membrane potential for SL neurons. **d** Phase-lag curves for SL neurons (mean ± SEM). **e** Quantification of phase-lag at 6 Hz in SL neurons. **f** Impedance profiles of AM neurons (mean ± SEM). **g** Quantification of *f_R_* as a function of membrane potential for AM neurons. **h** Phase-lag curves for AM neurons (mean ± SEM). **i** Quantification of phase-lag at 6 Hz (Φ_6Hz_) in AM neurons. **j** Scatter plot of *f_R_* vs. *R_in_* with the pooled data from SL and AM neurons. Colors indicate the voltage range of SL (circles, n=3x11) and AM (triangles, n=3x8) neurons. **k** Scatter plot of the same data of (j) transformed to logarithmic value. Labels are identical as (j). **l** Scatter plot of Φ_6Hz_ vs. *R_in_* from the pooled data of SL and AM neurons. Labels are identical as (j). All neurons were recorded at three voltage ranges. One way ANOVA, *=P<0.05,**=P < 0.01 and ***=P<0.001 and ****=P<0.0001.

To explore the changes in *f_R_* and phase-lag associated to the depolarization-driven increase in *R_in_*, we built scatter plots with the pooled data of *f_R_* and Φ_6Hz_ vs. *R_in_* from SL and AM neurons. *R_in_* ranged from 10 to ~100 MΩ in SL neurons and from 100 to ~350 MΩ in AM neurons, filling the *R_in_* range from 10 to 350 MΩ almost completely (Fig. 2j). *f_R_*s inversely correlate with *R_in_*, spanning the entire theta range when both data sets are taken together. The distribution of *f_R_* starts at 12 Hz for low-*R*_in_ SL neurons recorded at hyperpolarized potentials and decreases continuously to ~2 Hz for high-*R_in_* AM neurons recorded at depolarized potentials (Fig 2j). The distribution of *f_R_* and *R_in_* values, strongly suggest a multiplicative relationship, as described above (Fig. 1). The log-log graph exhibits an even higher linear correlation, with a Pearson’s coefficient of -0.92 and a slope of -0.43 (Fig. 2k, P<0.001). These results agree with the *R_in_-f_R_* relationship between different cell types, and show that individual neurons are able to modify their *f_R_* in coherence with changes in *R_in_*.

**Table 3.**
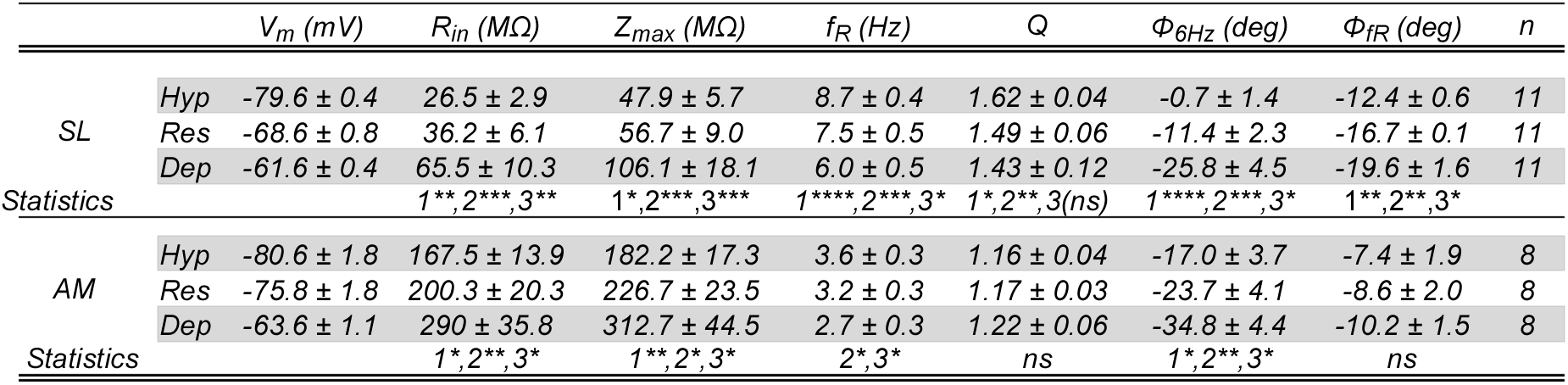
Intrinsic electrophysiological parameters for stellate (SL) and cortical amygdala neurons (AM) characterized at hyperpolarized (Hyp), near-resting (Res) and depolarized (Dep) potentials. Mean ± sem. Pairs for statistical comparison: 1 Hyp-Res, 2 Hyp-Dep and 3 Res-Dep. * P<0.05, **P<0.01, ***P<0.001, ****P<0.0001.

The scatter plot of Φ_6Hz_ against *R_in_* shows that cells increment their phase-lag during the depolarization-driven increase in *R_in_,* with both cell types spanning the whole range of 0 to -50 deg (Fig. 2l). This increase is dictated by linear correlations of different slope for each cell type (Pearson coefficient of -0.91 and -0.71 and slope of -0.45 deg/MΩ and -0.11 deg/MΩ for SL and AM neurons, respectively). Interestingly, despite having different *R_in_* ranges and phase-lag curves, both cell types can vary their Φ_6Hz_ along a similar range.

The question that arises is whether *R_in_* effectively has a causal role in the modulation of resonance. In single cells, *f_R_* is proportional to the level of activation of *I_h_* (Narayanan & Johnston, 2007), therefore it is expected that depolarization will deactivate a fraction of *I_h_* and reduce *f_R_.* Here, by definition, *R_in_* includes both, the passive and active electrophysiological properties; a decrease in *I_h_* with depolarization will also cause an increment in *R_in_* which, according to our results, will also be accompanied by a reduction of *f_R_*. Therefore, is not clear whether the changes observed in *f_R_* under the depolarization-driven increase in *R_in_* (Fig. 2) are due to changes in *I*_h_, *R_in_* or both, impeding a clean evaluation of the *f_R_-R_in_* relation.

To evaluate the causal role of *f_R_* modifications governed by *R_in_,* we designed a protocol to control *R_in_* while keeping unaltered the contribution of the voltage-dependent conductances involved in resonance. The observation that *f_R_* and phase-lag are modified when a neuron undergoes changes in *R_in_* with depolarization opens the possibility that other sources of *R_in_* variations, like synaptic activity, can operate as modulators of resonance. Both questions will be addressed in the following series of experiments.

### Modulation of the frequency preference and phase-lag by controlled changes in the input resistance

To explore additional evidence that physiological changes in *R_in_* can by themselves modulate the resonance we used the dynamic-clamp technique, to add or subtract virtual membrane conductances to the recorded neuron (Dorval et al., 2001). This virtual conductance is used to stimulate the cell, mimicking the opening (reducing *R_in_*) or closing (incrementing *R_in_*) of ion channels, thus closely representing a physiological condition, unlike the oversimplified traditional procedure of injecting current (Dorval et al., 2001). We performed these experiments in HP neurons at -80 mV, because at this voltage their *R_in_* values (~70 MΩ) are associated to *f_R_s* and phase-lags at the middle of the dynamic range, facilitating the detection of positive and negative changes (see Fig. 1).

To recreate a steady reduction (*-R_in_*) or increase (+R_in_) in *R_in_* we injected a dynamic current calculated as a constant leak conductance, *G_Leak_,* of positive or negative value, respectively (Fig. 3a,b. For more details see Methods). By manipulating the somatic conductance we aimed to reduce or increase the *R_in_* of HP neurons in order to match the average values of SL and AM cells, as examples of low and high *R_in_* cells, respectively. Using this criterion, we generated a *-R_in_* condition with an average reduction of ~45% (from 63.8 ± 5.4 MΩ to 35.4 ± 4.8 MΩ; *G*_Leak_=16.6 ± 2.9 nS, n=10, P < 0.05) and a ~100% increment for a *+R_in_* condition (from ± 4.7 MΩ to 151.8 ±16.7 MΩ; *G*_Leak_=-6.1 ± 0.8 nS, P<0.05, Fig. 3c). *R_m_* values of both control groups were not different (P = 0.13). These results demonstrate that our experimental procedure enabled us to change *R_in_* over a wide physiological range in different cell types. We manually adjusted the membrane potential to -80 mV and recorded the voltage responses to ZAP stimuli (Fig. 3d, black traces). Then, we reduced (-*R*_in_, blue traces) or increased (+*R*_in_, red traces) *R_in_* with the dynamic clamp and recorded the voltage responses to oscillatory stimulation at the new level of somatic conductance. In 4 out of 16 experiments we explored both conditions in the same cell, while in the other experiments we recorded only one of the two conditions and their respective controls. To keep constant the contribution of active properties at different *R_in_* values, we measured resonance systematically by adjusting the amplitude of the ZAP current to produce ~ 5 mV peak-to-peak oscillations (measured at 4 Hz, Fig. 3d). As expected, the manipulation of the somatic conductance was accompanied by changes in the impedance profile. In the *-R_in_* condition the impedance curve fell, decreasing in 50% at *Z_Max_* (from 90 ± 10.2 MΩ to 44.1 ± 6.2 MΩ). In contrast, in the *+R_in_* condition the impedance curves raised (Fig. 3e), reaching ~150% growth at *Z_Max_* (from 89.4 ± 5.1 MΩ to 221.6 ± 29.7 MΩ, Fig. 3f. P<0.01).

**Figure 3.**
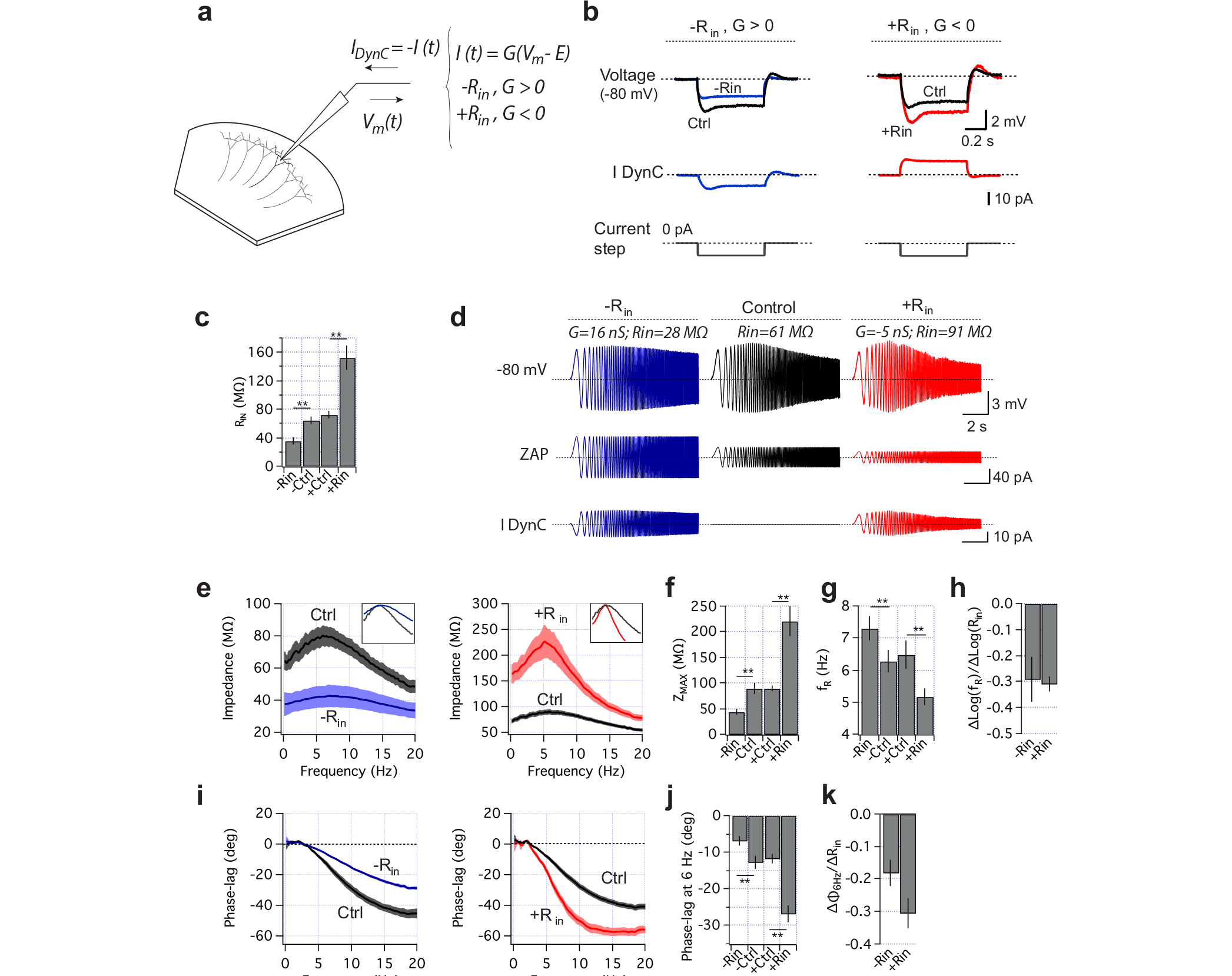
Changes in input resistance modulate frequency preference of CA1 hippocampal neurons. **a** The manipulation of *R_in_* was performed by means of the dynamic-clamp, as shown in the scheme. The neuron *R_in_* was decreased (*-R_in_*) or increased (+*R_in_*) injecting a virtual negative (+*G_Leak_*) or positive (*-G_Leak_*) somatic conductance as an external dynamic current (*I*_DynC_). **b** Dynamic-clamp experiment showing the attenuation (-*R_in_*) or amplification (+R_in_) of voltage responses revealed by the same 0.4 s and 50 pA hyperpolarizing pulse (gray). At-*R_in_* condition, *I_DynC_* acquires negative sign (blue trace), hyperpolarizing the neuron and attenuating voltage response. At +*R_in_*, *I_DynC_* is positive (red trace), depolarizes the cells and amplifies the voltage deflections. **c** Quantification of Rin manipulations as pairs, Control (-Ctrl) vs.-R_in_, Control (+Ctrl) vs. +*R*_in_. *R_in_* was measured with a squared pulse (as in b), but the amplitude was adjusted to evoke a ~4 mV voltage deflection in all conditions. The rationale to guide the magnitude of *R_in_* change was to attain a reduction of 50% (*G_Leak_* = 16.6 ± 2.9 nS, n=10) or an increase in ~100% (*G_Leak_* = -6.1 ± 0.8 nS, n=10). **d** Representative experiment showing oscillatory responses at different *R_in_* values; for simplicity we show an experiment in which both conditions were explored in the same cell. ZAP stimulation was applied at control (black), *-R_in_* (blue) or *+R_in_* (red) conditions. *I_DynC_* is the dynamic current injected by the dynamic-clamp. **e** Impedance profile curves (mean ± SEM) for *-R_in_* and *+R_in_* conditions, with their respective curves in the control condition. Note the difference in the scale in the impedance axes. The inset shows normalized curves. **e** Peak impedance for-*R_in_* and + *R_in_* conditions. **g** *f_R_* for *-R_in_* and +*R*_in_ conditions. **h** Ratio of Δlog (*f_R_*) and Δlog (*R*_in_) for *-R_in_* and +*R*_in_ conditions. **i** Phase-lag curves for *-R_in_* and + *R_in_* conditions, with their respective curves at control condition. **j** Phase-lag at 6Hz (Φ_6Hz_) for-*R*_in_ and *+R_in_* conditions. **k** Δ**Φ**_6Hz_/Δ *R_in_* for *-R_in_* and *+R_in_* conditions. **= P < 0.01, Paired Student T test.

The changes in peak impedance were accompanied by modifications in *f_R_.* In agreement with the results presented above, the reduction of *R_in_* involved an increase in *f_R_* of ~1 Hz (from 6.3 ± 0.4 to 7.3 ± 0.4 Hz, Fig. 3g), while an increase in *R_in_* was associated to a *f_R_* decrease of ~1.3 Hz (6.5 ± 0.4 Hz vs. 5.2 ± 0.3 Hz, Fig. 3g, P<0.01). To evaluate the degree of the resonance modulation, we calculated the ratio of the changes in the logarithms of *f_R_* and *R_in_* (Δlog(*f_R_*)/Δlog(*R_in_*) in each condition, obtaining values of -0.29 ± 0.09 and -0.31 ± 0.03 for *-R_in_* and +R_in_ conditions, respectively (Fig. 3h). These values are similar to the slopes of the linear regression obtained for log*f_R_*)-log(*R*_in_) when the source of *R_in_* variation was the heterogeneity of SL, HP and AM neurons (Fig 1k), or the depolarization-driven increase in *R_in_* in SL and AM neurons (Fig. 2k). This demonstrates that by modifying *R_in_* through changing only the passive somatic conductance it is possible to induce comparable variations in the frequency preference of the neurons.

*R_in_* manipulation also modified the phase-lag curve. The reduction of *R_in_* produced a frequency-dependent reduction in phase-lag that was traduced into a ~50% drop in Φ_6Hz_ (from -12.8± 1.7 deg to -6.8 ± 1.3 deg, Fig. 3i,j). For +*R_in_* the phase-lag increased, displaying a slower response at frequencies above 2 Hz (Fig. 3i), with a ~130% increase in lag (Φ_6Hz_ changes from -11.7 ± 1.3 deg to -26.9 ± 2.3 deg; Fig. 3j. P<0.005). These variations give a rate of change in Φ_6Hz_ (ΔΦ_6Hz_/Δ*R_in_*) of -0.18 ± 0.04 deg/MΩ and -0.31 ± 0.05 deg/MΩ in *-R_in_* and +*R_in_* conditions, respectively (Fig. 3i), between the values obtained for SL and AM neurons (Fig. 2l).

Taken together, these results indicate that by modifying exclusively the passive somatic conductance involved in setting *R_in_* it is possible to modulate resonance in the same ranges observed in figures 1 and 2, supporting a causal role of *R_in_* in setting and modulating the frequency preference and the phase-lag. Moreover, these observations suggest that individual neurons may adjust the frequency preference and the temporal response when *R_in_* varies, as it occurs upon intense synaptic input.

### Subthreshold modulation of the frequency preference and phase-lag is translated into changes in the frequency and timing of spikes

A critical question is whether the subthreshold modulation of the frequency preference is translated into a spiking regime, influencing neural activity. We tested this possibility by applying ZAP stimuli at perithreshold potentials and evaluating if changes in *R_in_* modified the spike firing probability and timing. These experiments were conducted in AM neurons because their high *R_in_* and low *f_R_* conditions offer a wide range to reduce *R_in_* and facilitates the detection of increases in firing frequency.

As in the previous experimental set, here we decremented *R_in_* by recreating a constant leak conductance with the dynamic-clamp. As expected, this manipulation decreased the voltage responses, with the consequent reduction in the excitability of the cell, but without altering its ability to fire action potentials (Fig. 4a). In these experiments we set the *-R_in_* condition by lowering *R_in_* in ~25 % (from 211 ± 27 MΩ to 163 ± 21 MΩ, with *G_Leak_=1.8* ± 0.46 nS, n=8. Fig. 4a). The neurons were depolarized to -60 mV and the amplitude of the oscillatory current was adjusted to reveal the preferential firing frequency. The stimuli consisted of a series of sinusoidal current pulses of constant amplitude with discrete increments in the frequency between 0.5 and 14 Hz (Fig. 4b, blue traces). A representative example of an AM neuron response is shown in Fig. 4b. (black trace). As expected, this neuron fired at frequencies of ~2-4 Hz, with no action potentials at higher or lower frequencies. Interestingly, in the *-R_in_* condition the neuron changed its firing preference (red trace) to higher frequencies (6-8 Hz), with no action potentials in lower range (2-4 Hz; Fig. 4c). To quantify the change in firing probability, we computed the cumulative firing probability (cumulative probability density function, CPDF) vs. stimulus frequency (Fig. 4d; see Methods) for all recorded neurons. These curves of the firing probability as a function of frequency show the distribution of selective firing frequencies from zero to one (Fig. 4d). AM neurons display a CPDF that rises from 0 at 2 Hz and reach a fractional probability of 1 at 8 Hz, limiting the firing probability to that frequency range (Fig. 4d). However, in the *-R_in_* condition the CPDF rises with a small probability at 2 Hz and saturates at 10 Hz, presenting lower firing probabilities than the control curve between 2 and 6 Hz (Fig. 4d, P<0.015). These differences indicate a displacement in the firing probability towards higher frequencies when *R_in_* is reduced. The comparison of the frequencies at which the CPDF curves reach a probability of 0.5 (F_P0 5_) reveals that the reduction of *R_in_* produces a ~2 Hz rightward shift in the firing probability curve (from 3.8 ± 0.7 Hz to 6.0 ± 0.8 Hz; P < 0.01, n=8, Fig. 4e). To discard that this firing modulation could have been generated by other factors that may also produce a reduction in *R_in_* and not by the *R_in_* change directly, we compared the spike threshold, the voltage at which a spike is generated (perithreshold potential) and the peak of the hyperpolarization induced with the ZAP protocol (hyperpolarized potentials) for each spike fired in both conditions (control and *-R_in_,* Fig. 4f). Neither of these parameters was different, suggesting that the voltage trajectory during oscillations and the activation of voltage-sensitive conductances was equivalent in control and *-R_in_* conditions. Therefore, the changes in firing frequency can be attributed exclusively to the changes in *f_R_* induced by the increase in somatic conductance. To evaluate if the modulation of the phase-lag influenced the timing of spikes fired, we quantified the phase-shift of each spike. We examined the delay of each spike relative to the peak of the sinusoidal current in control and *-R_in_* conditions (see Methods). When the oscillatory current reached ~4-8 Hz, the AM neurons fired action potentials with average delays of -13 to -18 deg, the same range observed for the subthreshold phase-lag (Fig. 4g, black traces). We found that in the *-R_in_* condition the neurons consistently fired action potentials with a reduced delay, at all tested frequencies (Fig. 4g, h). Note that at 4 Hz the reduction in the delay even caused that the spikes were fired earlier than the peaks of oscillatory current (Fig. 4g, h; vertical dotted line indicates the peak of oscillatory current used as the reference 0 deg). This reduction in the phase-shift is traduced into an average phase advance in spiking in 22, 18 and 13 deg for 4, 6 and 8 Hz, respectively (Fig. 4i, top trace). In the time domain, this manipulation gives an average advance in spike timing of 15.4, 8.4 and 4.6 ms for 4, 6 and 8 Hz, respectively (Fig. 4i, bottom trace). This result supports the notion that the modulation of the phase-lag associated to changes in *R_in_* at subthreshold potentials is also manifested in the spiking regime.

**Figure 4.**
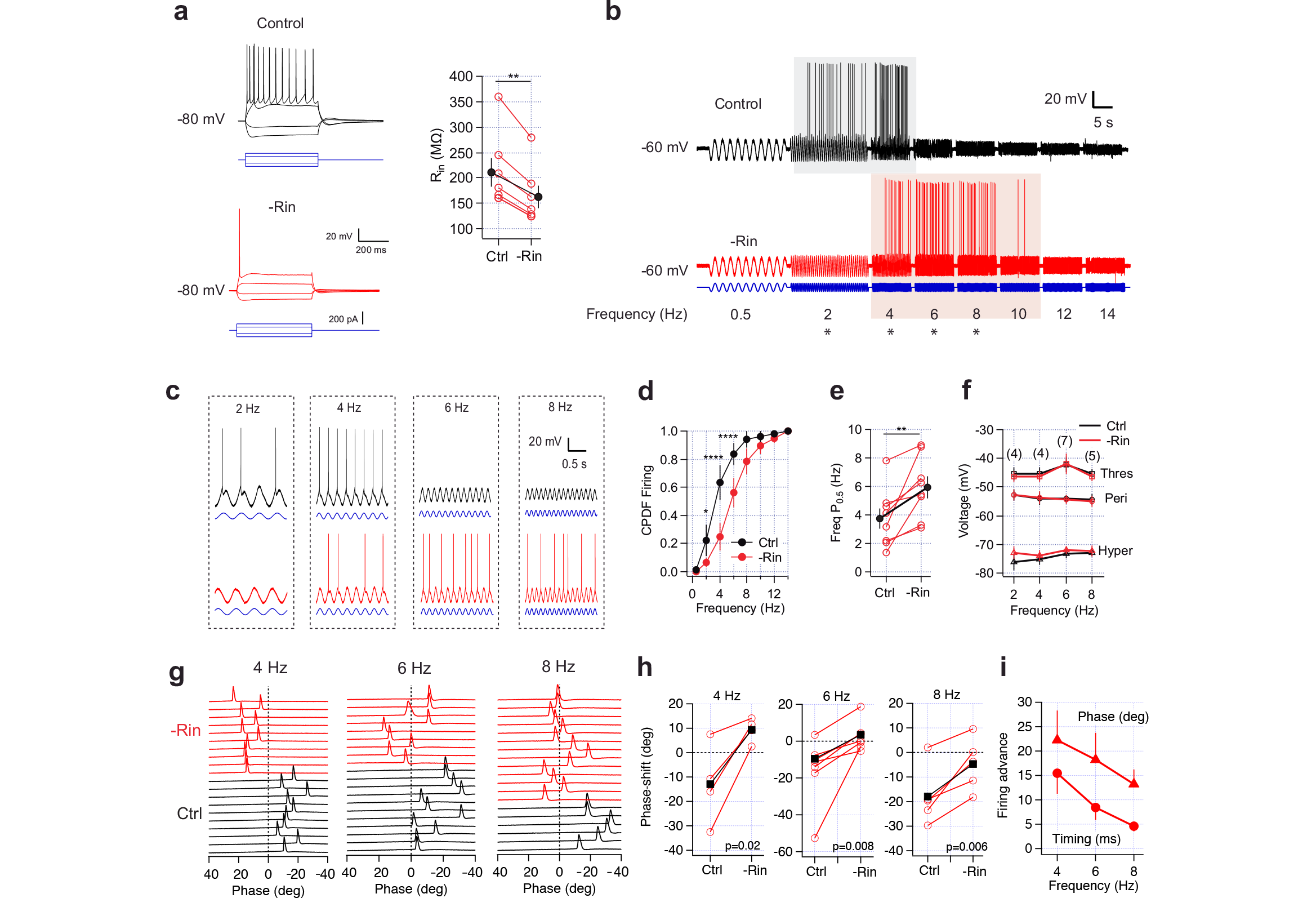
*R_in_*-driven modulation of subthreshold frequency preference is translated to spiking regime.

**a** voltage response of an AM neuron in control (black) and after a 30% reduction in *R_in_* (red, from 153 to 94 MΩ) adding a virtual somatic conductance by dynamic clamp. Current pulses are the same in both conditions. The plot shows paired measurement (empty circles) of *R_in_* drop in a set of AM neurons (n=8), filled circles are mean ± sem. **b** Voltage response of a resonant AM neuron under stimulation with an oscillatory current (blue) at control (black) and *-R_in_* (red) conditions. Stimulation protocol consisted in steps of discrete frequency at: 0.5, 2, 4, 6, 8, 10, 12 and 14 Hz (each frequency was applied during 10 s, with the exception of 0.5 and 2 Hz that lasted 20 s in order to sample more periods). Colored area shows the firing frequency range at each condition. **c** Magnification of recordings shown in b (*). **d** Cumulative probability density function (CPDF) of firing probability curves at control and *-R_in_* conditions (n=8, see Methods and main text for details). **e** Paired measurement of the F(P_05_) (empty circles) at both conditions, filled circle are the mean ± sem. **f** Average curves for spike threshold (Thres), perithreshold membrane potential (average of 20 ms window before spike threshold) and the peak of hyperpolarization during ZAP stimulation, at control and *-R_in_* conditions. Numbers above each curve indicate cells that fired action potentials at both conditions. **g** Representative recordings showing the phase-lag of spikes fired at control (black) and *-R_in_* (red) condition. Each perturbation in voltage traces is a single spike. The phase was calculated using the positive peak of sinusoidal current stimuli as a reference. **h** Paired quantification of phase shift in control and *-R_in_* condition (empty circles), filled circles are average values. **i** Firing advance quantified as the difference in phase-lag between control and *-R_in_* condition. Phase-lag is in deg, and timing in ms. *=P<0.05, **=P<0.01, ****=P<0.0001.

### A resonant current and a leak conductance are sufficient to account for the setting and modulation of the frequency preference and temporal response

We used computer simulations to evaluate if a minimal conductance-based model containing only the hyperpolarization-activated conductance (*G_h_*), a passive leak conductance (*G_Leak_*, the reciprocal of *R_m_*) and a membrane capacitance (*C_m_*), can account for our experimental observations.

We first aimed to reproduce the differences in resonant properties of SL, HP and AM cells. Our strategy was to set the value of *C_m_* according to experimental measurements and then adjust the magnitudes of *G_h_* and *G_Leak_* for each cell type to reproduce the average behavior observed under ZAP stimulation at -80 mV (i.e., *R_in_, *f_R_** and phase. Fig. 5a, dotted box). Using this minimal model, it was possible to reproduce the characteristic voltage response of each cell type (Fig. 5a, dotted box. See details in Methods). Moreover, varying C_m_, *G_h_* and *G_Leak_* within physiological values, it was possible to reproduce an *f_R_* value inside the theta range and a proper phase-lag, in a way that mimics the experimental results. We then used the three model neurons to explore the effect of *R_in_* variations caused by changes in the passive conductance *G_Leak_.* As in our dynamic-clamp experiments (Fig. 3), we explored a reduction (-*R*_in_) or an increase (+R_in_) of the input resistance by manipulating *G_Leak_* gradually between 1 and 90 nS. This manipulation varied the *R_in_* of the model cells over the whole range, from 10 to 310 MΩ. Each simulated neuron changed their frequency preference and phase-lag as a result of *G_Leak_* modifications. The reduction of *R_in_* produced an increase in *f_R_* and a reduction in Φ_6Hz_ in the three cell types, with opposite effects when *R_in_* is increased (Fig. 5a, arrows indicate the *f_R_*). Moreover, the *f_R_* vs. *R_in_* relationship obtained for each simulated cell type displays the same power-law distribution observed in our experimental data (Fig. 5b, filled circles show values at *G_Leak_* control value, representing the average behavior of each cell type as in Table 2). Note that these simulations represent the range of modulation of a single neuron when *R_in_* was varied with slight increments in *G_Leak_.* In agreement with the experimental measurement, the log(*f_R_*)-log(*R*_in_) plot yields nearly linear curves in the three model cells, described by a linear fit with slopes of -0.29 and -0.31 for the SL and HP models, respectively (Fig. 5c). In the case of the AM model, the log-log curve presents two regions of different slopes, of -0.25 below 2.0 Hz (similar to the HP model) and -0.57 above 2.0 Hz (Fig. 5c, dotted line above green line).

**Figure 5.**
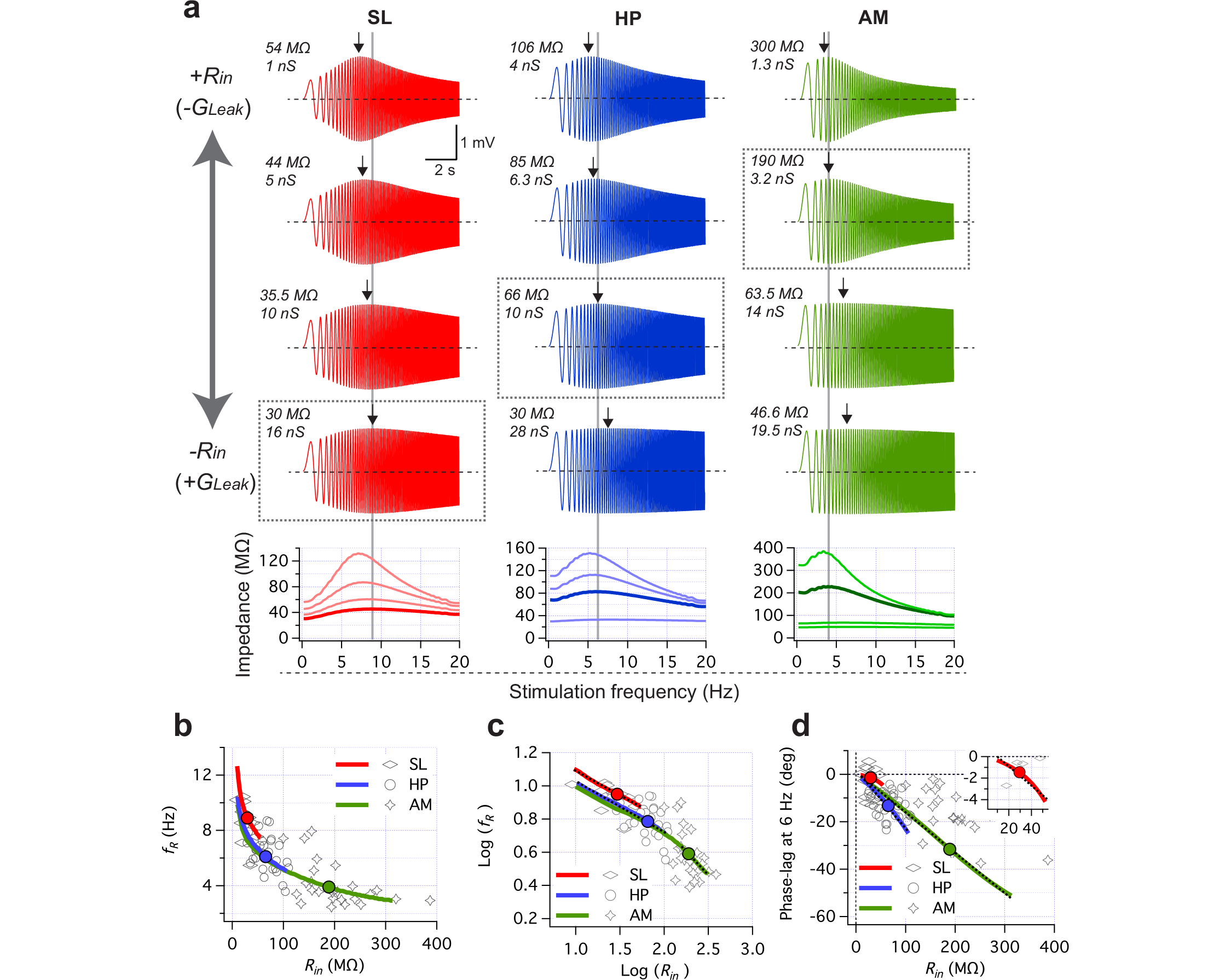
Minimal computer model reproduce the setting and modulation of frequency preference and phase-lag according to *R_in_* range.

**a**. A representative SL, HP and AM neurons were simulated using a minimal resonant model (see Methods). Starting with a *G_Leak_* that reproduced the average *f_R_, R_in_* and Phase-lag of each cell type (i.e. control conditions, dotted box), we explored voltage responses ranging *G_Leak_* from ~1 nS (*+R_in_*) to 80 nS (*-R_in_*) while maintaining constant *I_h_* and *C_m_* (4 conditions are shown for each cell type, *R_in_* and *G_Leak_* are reported at the top-left of each voltage trace). The impedance profiles are shown at the bottom plot, with the darker curve corresponding to the control condition. Grey lines show the *f_R_* at control condition, while arrows show the *f_R_* at each voltage trace. The amplitude of oscillatory current was adjusted to produce voltage responses of ~2 mV peak-to-peak while DC current was injected to hyperpolarize to -80 mV. **b**. Scatter plot of *f_R_* vs. *R_in_* obtained with the exploration of *G_Leak_* in the three cell types (colored lines), with filled circles showing values at control condition. Empty figures show experimental data. **c**. Same as b but plotting log(*f_R_*) vs. log(*R*_*in*_). Dotted black lines are the linear fit of each curve, with a slope of -0.29 (SL), -0.31 (HP) and -0.57 (AM, note this fit is between 2.2 and 2.5. See main text). **d**. Scatter plot of Φ _6Hz_ vs. *R_in_* obtained from *G_Leak_* exploration (colored lines), empty figures are experimental data. The inset shows a zoom to visualize SL response.

The exploration of different *R_in_* values in our model cells also reproduced the modification of the phase-lag, increasing the Φ_6Hz_ proportionally to *R_in_,* with a distribution that matches the experimental values (Fig. 5d). The rate of change was different for each model neuron, with slopes of -0.09, -0.25 and -0.17 deg/MΩ for SL, HP and AM models, respectively (Fig. 5d, dotted lines).

Overall, the computer simulations demonstrate that only one resonant conductance (in this case the h current) is sufficient to provide a mechanism of frequency preference that is set and dynamically modulated by the magnitude of the membrane conductance.

To illustrate the mechanism by which *R_in_* influences the *f_R_* and phase-lag of neurons, we used a conductance-based model to qualitatively analyze the independent contribution of the low-pass and high-pass filters in setting the *f_R_* and the phase-lag of the response after a *R_in_* change. Figure 6a shows the decomposition of a resonant impedance profile into 4 components: i) the low-pass filter produced by *R_m_* and *C_m_* (blue), ii) the high-pass filter produced by the resonant current (I_h_, black), iii) the resultant band-pass filter (dark orange) and iv) the impedance of the trivial case in which only *R_m_* is considered (i.e., in the absence of *C_m_* and inductive current) as the inverse of total membrane conductance (gray). This resistive impedance sets the value of the membrane resistance (*R_m_* = ~90 MΩ) with slight frequency dependence, determining the maximal values for the impedance at both, the low and high-pass filters. Therefore, the bandpass filter that accounts for the frequency preference in the theta range results from the meeting point of the low and high-pass filters, defining a set-point for *f_R_* (red circle).

The simplest scenario to describe mechanistically the modulation of the *f_R_* and the phase-lag of the response is the variation of *R_in_* due to changes in a leak conductance (as in Figs. 3 and 5). A reduction in *G_Leak_* (increase in *R_m_* and thus in *R_in_*) raises the maximal impedance for both, low and high-pass filters (+R_in_ condition, Fig. 6, green arrows). Since the steepness of the low-pass filter is set by its cut-off frequency (f_c_=1/2π*R_m_*C_m_), the increase in *R_m_* reduces this value and increases its steepness (blue circle and green arrow). On the other hand, the cut-off frequency of the high-pass filter is independent of *R_m_*, thus its frequency dependence remains constant while its maximal impedance is updated to the new value of *R_m_.* The modifications in the low-pass filter (increase in maximal value and steepness) displace this filter towards low frequencies, shifting the meeting point with the high-pass filter towards low frequencies and reducing *f_R_* (Fig. 6a). In case of a reduction of *R_in_* by an increase in *G_Leak_*, the modifications occur in the other direction. The increases in *G_Leak_* reduce *R_m_* (and *R_in_*) and rises the cut-off frequency of the low-pass filter, reducing its steepness, displacing the meeting point with the high-pass filter towards higher frequencies and increasing *f_R_* (Fig. 6a, *-R_in_* condition red arrows).

**Figure 6.**
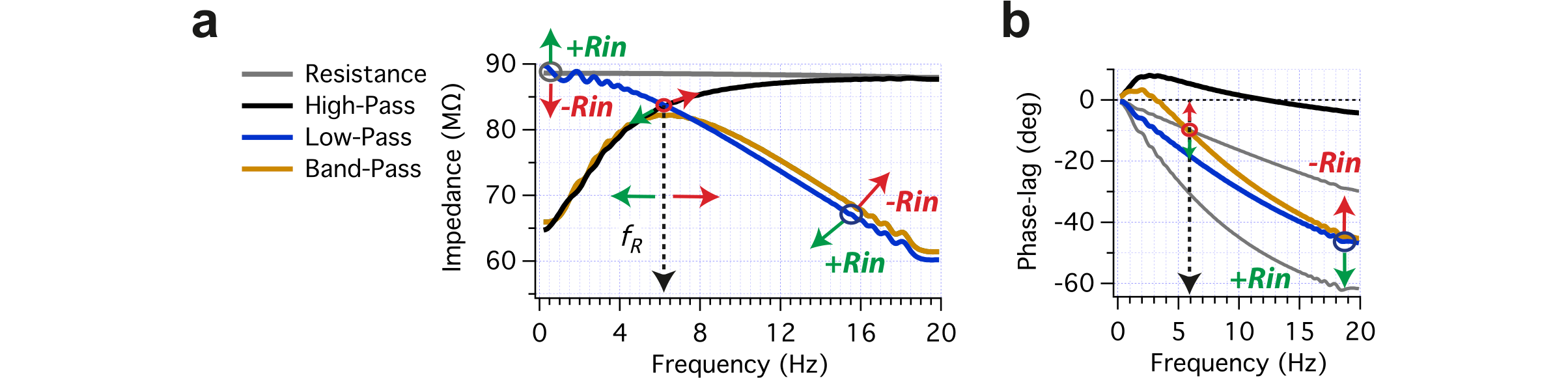
Mechanism behind the modulation of frequency preference and phase-lag by changes in *R_in_.* **a.** Independent impedance analysis of resistive (grey), high-pass (black), low-pass (blue) and band-pass filters (dark yellow) of a resonant neuron. A model cell consisting in a capacitance (100 pF), a leak conductance (*G_Leak_*=10 nS) and an *h* conductance (*G_h_*= 3 nS) was used to investigate the impedance profile of each component. Each simulation was performed maintaining the cell at -80 mV and stimulating with the ZAP current (0-20 Hz in 10 s interval) producing a maximal peak-to-peak response of 2 mV. **b.** Phase-lag curves obtained from the impedance analysis described in a. Arrows indicate the direction of curve changes when *R_in_* is increased (*+R_in_*, green) or reduced (*-R_in_*, red).

In the case of the phase response, the observation of the resistive, capacitive and inductive components separately allows an easier understanding of the role of total conductance in the control of the temporal properties of the response (Fig. 6b). The phase response associated to the low-pass filter (*R_m_* and *C*_m_) shows a monotonic increase in lag along the frequencies of interest (Fig. 6d, blue). In contrast, the phase response of the high-pass filter shows positive lead with a peak near 3 Hz, which gradually descends and turns to negative lags around 10 Hz (black). When the complete model is evaluated, the phase-response is the sum of the other two curves (high and low-pass), displaying a positive phase lead below 3 Hz due to the inductive effect, and a shift to negative lags at higher frequencies (dark orange trace). Following the previous example, a reduction in *G_Leak_* increases *R_m_* (and *R_in_*) with a consequent increase in the phase-lag due to the low-pass filter (Fig. 6b, green arrows and bottom gray curve). Since the phase response of the high-pass filter is not altered, the sum of both curves yields a phase response with higher lag, increasing the lag at 6 Hz (dotted arrow). When *G_Leak_* is raised, the low-pass filter reduces the lag, with a consequent reduction in the lag of the final curve (red arrows). This dependence of the phase-lag curve on *R_m_* explains the modulation of the phase observed experimentally.

Taken together, these results show that the *f_R_* and phase-lag of neurons are dynamic parameters that vary according to *R_in_*, serving as a tuning mechanism to modulate the frequency preference inside theta range, and to adjust the timing of action potential generation into a millisecond scale.

## Discussion

In the present study we investigated the role of the input resistance in setting the frequency of preference and phase-lag upon oscillatory stimulation in cortical neurons. We found that the input resistance determines the *f_R_* following a power law, in which *R_in_* in the physiological range sets the *f_R_*s spanning the theta frequency range. Moreover, the phase-lag of the responses critically depends on *R_in_.* This dependence on *R_in_* has two major implications: 1) Cell types within different *R_in_* ranges will display distinct frequency preference and phase-lag, following a stereotyped scheme, and 2) Changes in *R_in_,* as those expected to occur instantaneously in physiological conditions, modulate the frequency preference and phase-lag along the theta range. Additionally, we show that this modulation can be translated into a firing regime, modifying the spike firing frequency and timing in a way that might be relevant for the network activity (Lisman and Jensen, 2013; Ratté et al., 2013).

### *R_in_* sets and modulates the preferred frequency of cortical neurons

The conclusions summarized above are based on experimental measurements and computer simulations. The first evidence is derived from a comparative analysis across different resonant cell types that allowed us to characterize, under the same experimental conditions, their resonance properties along the range of *R_in_* normally observed in cortical neurons. This comparison revealed a power law tendency between *f_R_* and *R_in_*, with *R_in_* values spanning the whole theta range. When transformed into a log-log plot, the data from the three cell types present a linear relationship, thus offering a simple way to describe and quantify the *f_R_-R_in_* link. Therefore, we propose that resonant neurons with different ranges of *R_in_* will show different set-points for *f_R_* in the theta range, a value that can be described by a simple mathematical relationship.

A second line of evidence confirmed that variations in the *R_in_* of individual neurons are accompanied by modifications in their frequency preference and temporal responses (Fig. 2). Considering the increase in *R_in_* that occurs when the neurons are depolarized (Surges *et al*., 2004; Economo *et al*., 2014; Yamada-hanff & Bean, 2015; Ceballos *et al*., 2017), we measured the voltage response of SL and AM neurons at three different *R_in_* values. The variation of *f_R_* with respect to *R_in_* exhibited an even higher degree of correlation. The slope of the log-log linear distribution of *R_in_* and *f_R_* was in close agreement with the value obtained for different cell types (Fig. 1). This result shows that the *R_in_-f_R_* relationship also applies for *R_in_* variations in individual neurons within physiological ranges and that these changes may dynamically modulate the *f_R_* along the theta range.

To confirm the causal role of *R_in_* in this modulation we changed the *R_in_* by modifying only its passive component, increasing or decreasing somatic conductance by means of dynamic-clamp (Dorval *et al*., 2001). This technique makes possible to maintain unchanged the active and preexistent passive components and observe the resulting voltage response. In this unique experimental condition, we reproduced the changes in *f_R_* due to variations in *R_in_*. Moreover, we further corroborate with computer simulations the strength of the *R_in_-f_R_* relationship, and show that the presence of one resonant current (in this case *I_h_*) and a leak conductance are sufficient to fully reproduce the modulation of *f_R_* and phase-lag associated to changes in *R_in_* (Fig. 5).

Additionally, the use of computational models allowed us to evaluate the entire possible range of *f_R_* modulation as a function of *R_in_*. We found that a single neuron can change its *f_R_* along most of the theta range, regardless of the cell type. Considering the constrains of these minimal models, it is expected that the diverse set of voltage and ligand-gated ion channels present in real membranes would expand the range of *f_R_* modulation along the whole theta range. It can be noted, for example, that the neurons duplicate *R_in_* relative to hyperpolarized values near the spike threshold (due to the depolarization-driven increase in *R_in_*). Besides, the contribution of the active membrane properties and morphology (size and shape) of neuron will provide a structural constrain to set *R_in_* into a defined range at resting conditions. Moreover, considering the complex cell morphology and the heterogeneous expression of the ion channels implicated in the resonance (responsible of *I_h_* and *I_NaP_*), it is possible that *R_in_* changes can occur independently in different sites of the same neuron. This would produce an even more complex scenario, where *f_R_* would be modulated differentially in separate compartments of the neurons according to the local variations in membrane conductance.

It is important to mention that our experiments were performed under conditions that favored the stability of the recordings (-80 mV and reduced voltage oscillations), rather than maximizing the magnitude of the modulation. For example, as explained below, it is expected that at perithreshold potentials the modulation of the resonance will be higher, since the peak impedance can reach above 100 MΩ (Surges *et al*., 2004; Economo *et al*., 2014; Yamada-hanff & Bean, 2015; Ceballos *et al*., 2017; Vera *et al*., 2017) even in SL neurons (Table 3), providing a wider dynamic range to vary *R_in_* and modulate *f_R_* and phase-lag.

### *R_in_* sets and modulates subthreshold phase-lag

In contrast to the importance usually assigned to *f_R_,* the role of the phase-lag in processing the oscillatory inputs has received less attention (Hutcheon et al., 1996a; Hutcheon and Yarom, 2000; Richardson et al., 2003). What makes the modulation of the phase-lag in resonant neurons interesting is that it introduces a novel aspect of the frequency preference, since the timing of the spikes generated selective at the *f_R_* could be adjusted. The dependence of *f_R_* and phase-lag on *R_in_* implies that both parameters vary together, with higher *f_R_* being associated with lower lag. Subthreshold phase-lag has been shown to depend on *I_h_* in CA1 pyramidal neurons, and the proximal to distal gradient of the *I_h_* conductance along the dendrites controls the magnitude of the phase-lag (Narayanan & Johnston, 2008). The increased *G_h_* conductance at distal sites reduces the phase-lag of distal inputs, compensating for the delay associated to dendritic filtering and producing temporal synchrony between distal and proximal inputs (Vaidya & Johnston, 2013). One of the most interesting aspects of the resonance is that during an oscillatory activity the inductive properties of the membrane allow a zero or even a positive phase lead at a specific frequencies (Narayanan & Johnston, 2008; Rotstein, 2014).

Here we show that the modulation of the phase-lag in the subthreshold response is preserved in the spiking regime as a modulation in the delay of the spikes fired when the voltage goes suprathreshold, influencing directly the timing of the neuronal responses (Fig. 4). The proper control of spike timing is vital for the generation of synchronized activity between different brain circuits (Fuhrmann *et al*., 2002), suggesting a novel mechanism to precisely adjust the spike timing in the millisecond scale. According to theories of neural coding based on nested theta-gamma activity and neural synchrony, the control of the spike timing is crucial to properly coding and decoding neuronal information (Lisman and Jensen, 2013). Therefore, it can be speculated that neurons exploit the *R_in_*-Phase-lag relationship to adjust their temporal responsiveness in order to properly engage in coherent activity. Until now, a detailed measurement of phase-lag in cortical neurons as a function of *R_in_* and voltage had not been done. Our comparative study allowed the characterization of the subthreshold phase-lag curves for the different cell types and to measure their variation under *R_in_* changes, finding that maximal differences in phase-lag curves among the three cases occur in the theta range.

The phase response of the three cell types was very different when compared at 6 Hz, Φ_6Hz_ was close to 0 deg for SL neurons, increasing substantially in HP and AM neurons. These marked differences in the phase responses are reduced when compared at the *f_R_* of each cell. This similarity of Φ_fR_ among cells with different phase-lag curves is the result of an opposite effect of *R_in_* changes on *f_R_* and phase-lag. While a *R_in_* reduction increases *f_R_* and displaces Φ_fR_ toward more negative values of lag (the lag increases with frequency), it reduces the amplitude of the new phase-lag curve (low *R_in_* implies a reduced lag) in a way that compensates for phase-lag measured at *f_R_,* thus keeping relatively similar values (Tables 1, 2 and 3). Given this compensatory effect, it is reasonable to think that cortical neurons that oscillate near their *f_R_* will share similar phase-lags. Therefore, resonant neurons promote a constant delay in the transmission of electrical activity regardless of their individual *f_R_*s. Whether this intriguing phenomenon has some role in the processing of oscillatory activity has to be investigated.

### Modulation of the resonance is translated into spiking regime

Here we confirmed that subthreshold frequency preference is a property that can be translated into a selective firing at the preferred frequency, as previously described for neocortical neurons (Hutcheon *et al*., 1996a; Ulrich, 2002), SL neurons (Erchova *et al*., 2004), HP neurons (in this case relying on *I_M_* as a resonant current) (Vera *et al*., 2017) and AM neurons (Vera *et al*., 2014). More importantly, our results reveal that the modulation of *f_R_* and phase-lag is preserved during the spiking regime, generating a dynamic tuning of both, the firing frequency and the spike timing. Since this tuning mechanism relies on the modulation of basic electrophysiological properties (i.e. the subthreshold frequency preference and phase-lag), it is reasonable to propose that this is a general property exhibited by most cortical neurons.

### Intrinsic changes in *R_in_*: Self-generated modulation of resonance

The mechanism that we describe here points to the cell impedance as a key factor in the set-point of the frequency preference and phase-lag. Since *R_in_* will vary in cortical neurons expressing subthreshold voltage-dependent conductances (like *I_h_, I_M_* or *I*_NaP_)(Surges *et al*., 2004; Yamada-hanff & Bean, 2015; Ceballos *et al*., 2017; Vera *et al*., 2017), their resonant properties will depend on the membrane potential. At hyperpolarized potentials, *I_h_* will be fully activated, reaching a maximum in the steady-state conductance and a consequent minimum *R_in_.* At this point, the neurons will express their intrinsic highest values of *f_R_* and lowest phase-lag (fastest response). At depolarized potentials (above -70 mV), *I_h_* deactivation will reduce the steady-state conductance, increasing the impedance, shifting *f_R_* to lower values and increasing the phase-lag. Above -70 mV *I_NaP_* activates, producing a significant increase in *R_in_* (Ceballos *et al*., 2017; Vera *et al*., 2017), and above -60 mV *I_M_* begins to activate with a consequent reduction in *R_in_.* Given this combined effect of *I_h_* deactivation and *I_NaP_* and *I_M_* activation, the neurons are able to increase in 100% or more their *R_in_,* thus significantly reducing *f_R_*s and increasing Φ_6Hz_. In this context, *I_NaP_* might play a pivotal role in tuning of the frequency preference and temporal neuron response, raising the perithreshold *R_in_* and increasing the range for *f_R_* and phase-lag modulation.

### Changes in *R_in_* due to synaptic inputs: Extrinsic control of the frequency preference

In addition to *R_in_* changes resulting of intrinsic modifications, a major source of *R_in_* variation are synaptic inputs. Regardless of their excitatory or inhibitory nature, the activation of post-synaptic receptors increases the membrane conductance with a consequent reduction in *R_in_.* Moreover, it has been postulated that neurons that are part of active circuits *in vivo* receive a plethora of uncorrelated excitatory and inhibitory synaptic inputs that gives origin to drastic *R_in_* variations, that can drop to 10% of the values observed *in vitro* (reviewed in Destexhe et al., 2003). According to our results, such *R_in_* variations during intense synaptic activity would change the *f_R_* in ~2 Hz and reduce Φ_6Hz_, causing a shift of the output wave of more than 6 ms. Importantly, this mechanism of modulation enables upstream neurons of the network to strongly influence the frequency preference and temporal response of their target cells, controlling their *f_R_* and phase-lag through differential levels of synaptic activity.

Therefore, the *f_R_* of a cortical neuron in a specific moment is set along the theta range according to its *R_in_*, which in turn is dynamically adjusted by the interaction between the intrinsic and synaptic properties. The same occurs with the phase-lag. Whether cortical circuits effectively implement this mechanism of *f_R_* modulation to promote oscillatory activity in the theta range requires further investigation.

The dynamic adjustment of the frequency preference and phase-lag may contribute to engage cortical neurons with the time-varying theta activity of the network, providing a robust cellular mechanism to promote coordinated neuronal activity across separate brain regions.

## Acknowledgments

We thank Pablo Villar for To explore helpful discussions and Dr. Alain Marty for thoughtful comments on the manuscript.

Funded by the Chilean Fund for Science and Technology (FONDECYT) 1140700 (MS), 3150668 (JV) and 1140520 (JB).

